# Activation of WNT signaling restores the facial deficits in a zebrafish with defects in cholesterol metabolism

**DOI:** 10.1101/2020.02.14.949958

**Authors:** Victoria L. Castro, Nayeli G. Reyes-Nava, Brianna B. Sanchez, Cesar G. Gonzalez, Anita M. Quintana

## Abstract

**Background:** Inborn errors of cholesterol metabolism occur as a result of mutations in the cholesterol synthesis pathway (CSP). Although mutations in the CSP cause a multiple congenital anomaly syndrome, craniofacial abnormalities are a hallmark phenotype associated with these disorders. Previous studies have established that mutation of the zebrafish *hmgcs1* gene (Vu57 allele), which encodes the first enzyme in the CSP, causes defects in craniofacial development and abnormal neural crest cell (NCC) differentiation. However, the molecular mechanisms by which the products of the CSP disrupt NCC differentiation are not completely known. Cholesterol is known to regulate the activity of WNT signaling, an established regulator of NCC differentiation. We hypothesized that defects in cholesterol synthesis reduce WNT signaling, consequently resulting in abnormal craniofacial development.

**Methods:** To test our hypothesis we performed a combination of pharmaceutical inhibition, gene expression assays, and targeted rescue experiments to understand the function of CSP and WNT signaling during craniofacial development.

**Results:** We demonstrate reduced expression of *axin2*, a WNT downstream target gene in homozygous carriers of the Vu57 allele and in larvae treated with Ro-48-8071, which inhibits the synthesis of cholesterol. Moreover, activation of WNT signaling via treatment with a WNT agonist completely restored the craniofacial defects present in the Vu57 allele.

**Conclusions:** Collectively, these data suggest interplay between the CSP and WNT signaling during craniofacial development.

## Introduction

The cholesterol synthesis pathway (CSP) produces two unique lipid molecules, cholesterol, an essential membrane component and precursor for steroids (Craig & Malik, 2020), and isoprenoids, a class of lipids essential for cell signaling (Lange et al., 2000). Mutations that inhibit this pathway cause inborn errors of cholesterol synthesis. There are 8 different inborn errors of cholesterol metabolism, each caused by mutations in a different enzyme of the CSP (Porter, 2002). These mutations cause a multiple congenital anomaly syndrome, however, craniofacial abnormalities are a hallmark phenotype of these disorders (Porter, 2002). Multiple model systems have been developed in mice (Nwokoro et al., 2001) and zebrafish (Quintana et al., 2017, p. 1; Signore et al., 2016) to understand the mechanisms by which the products of the CSP regulate facial development.

The bone and cartilage of the face develops from neural crest cells (NCCs), a multipotent progenitor cell that arises from the dorsal end of the neural tube. A subset of NCCs, cranial NCCs (CNCC), migrate, proliferate, and differentiate into bone and cartilage structures (Bhatt et al., 2013). We and others have demonstrated that cholesterol is an essential mediator of CNCC differentiation and that the inhibition of cholesterol causes facial defects in zebrafish (Quintana et al., 2017; Signore et al., 2016). However, the molecular mechanisms underlying these differentiation defects are not completely known. Cholesterol is known to regulate cell signaling in the form of lipid rafts and via the modification of specific proteins (Sheng et al., 2012). For example, cholesterol modification of sonic hedgehog (SHH) is essential for activation (Koleva et al., 2015).

Activation of SHH is essential for proper facial development and therefore, in previous studies we hypothesized that cholesterol regulates facial development via activation of SHH. However, mutation of the CSP in developing zebrafish causes defects in facial development in the absence of abnormal SHH signaling (Quintana et al., 2017). These data strongly suggest that cholesterol regulates facial development in a SHH independent manner. Several other signaling cascades are essential for proper facial development, including FGF, BMP, WNT, and retinoic acid (Shimomura et al., 2019). Of these, cholesterol is required for activation of canonical WNT signaling via the PDZ domain in the disheveled protein (Sheng et al., 2012). Therefore, we hypothesized that mutation of the CSP causes facial abnormalities due to defects in WNT activation.

To test this hypothesis, we performed a combination of loss of function, pharmacological inhibition assays, and restoration experiments using the developing zebrafish. The inhibition of cholesterol synthesis was associated with a significant decrease in the expression of WNT activity (*axin2*) and the expression of *sox10*, a marker of NCC cells (Kelsh, 2006, p. 10). The activation of WNT using a pharmaceutical agonist completely restored the facial deficits present in zebrafish mutants of the CSP. Collectively, these data suggest that cholesterol regulates facial development in a WNT dependent manner.

## Materials and Methods

### Zebrafish Care/Embryo Collection

Embryos were obtained by mating male and female pairs of Tupfel long fin, *hmgcs1*^Vu57^ (Mathews et al., 2014), *Tg*(*sox10*:memRFP) (Kucenas et al., 2008), or *Tg*(*sox10*:tagRFP) (Blasky et al., 2014). Embryos were maintained in an incubator at 28°C and all experiments were performed at The University of Texas at El Paso according to approved guidelines from the Institutional Animal Care and Use Committee (IACUC) protocol 811689-5.

### Genotyping

Genotyping was performed as described in (Quintana, Hernandez, & Gonzalez, 2017) and (Matthews, et al., 2014). DNA was extracted from all samples using phenol:chloroform ethanol precipitation as described in (Quintana, Hernandez, & Gonzalez, 2017). Samples were collected either by clipping the fins of adult zebrafish, collecting eggs, or dissecting the embryos and using a small piece of tissue for DNA extraction. DNA was extracted in DNA extraction buffer (10mM Tris pH 8.2, 10mM EDTA, 200mM NaCl, 0.5% SDS, and 200ug/ml proteinase K) for three hours at 55°C. Polymerase chain reaction (PCR) was used to amplify the target sequence of the *hmgcs1* gene. PCR and restriction digest was performed as previously described in (Quintana, Hernandez, & Gonzalez, 2017) (Hernandez et al., 2019).

### RNA extraction and in situ hybridization

RNA isolation was performed with Trizol according to manufacturer’s protocol (Fisher). Total RNA (100-500ng) was reverse transcribed using the Verso cDNA synthesis kit (Fisher) and cDNA was used for quantitative real time PCR. Quantitative real time PCR (qPCR) was performed as described in (Quintana, Hernandez, & Gonzalez, 2017). Primers for qPCR are as follows: 1) *Axin2 fwd* 5’-CAACCAAGCACATCCATCAC-3’ and *rev* 5’-TCCATGTTCACCTCCTCTCC-3’, 2) *rpl fwd* 5’-TCCCAGCTGCTCTCAAGATT-3’ and *rev* 5’-TTCTTGGAATAGCGCAGCTT-3’, and 3) *Sox10 fwd* 5’-ACGCTACAGGTCAGAGTCAC-3’ and *rev* 5’-ATGTTGGCCATCACGTCATG-3’. Whole mount in *situ* hybridization was performed as previously described (Hernandez et al., 2019; Quintana et al., 2017; Thisse & Thisse, 2008). The *axin2* riboprobe was produced with forward primer 5’-TGCACTGCTCCTTACATTCG-3’ and reverse primer 5’-GCTCCAGGACAAGGCTACTG-3’.

### Drug Treatments

Wnt agonist-1 was dissolved in 100% DMSO and embryos were treated with a final concentration ranging from 0.1-1.0 μM. Concentrations were developed from recommendations previously described (Liu et al., 2005). Final concentration of DMSO was less than 0.01% in all treated samples and the vehicle control treatment. Wnt agonist-1 treatment was initiated at 30 hours post fertilization (hpf), consistent with the onset of facial defects (Quintana et al., 2017) and removed at 54hpf, for a total treatment time of 24 hours of treatment at 28°C. Treated embryos were maintained at 28°C until 4 days post fertilization (dpf). Treatment with Ro-48-8071 was performed as previously described (Hernandez et al., 2019; Quintana, Picchione, et al., 2014; Quintana et al., 2017).

### Staining and Imaging

Staining was performed as described in (Quintana, Geiger, et al., 2014). Briefly, embryos were harvested at 4 dpf and fixed in a 2% PFA and 7.4 pH PBS solution for 1 hour. After fixing, embryos were stained with Alcian Blue (Anatech Ltd., #862 5 mL 0.4% Alcian Blue in 70% EtOH, 5 mL Tris pH 7.5, 0.5 mL 1M MgCl2, 36 mL EtOH 80%) overnight at room temperature. Samples were washed with 80% EtOH/100mM Tris pH 7.5/10mM MgCl2 for 5 minutes, 50% EtOH/100mM Tris pH 7.5 for 5 minutes, and 25% EtOH/100mM Tris pH 7.5 for 5 minutes, each at room temperature. Pigment was removed by incubation in bleach buffer (100μL 30% H2O2, 25 μL 20% KOH, 875μL H2O) for 10 minutes at room temperature. Bleach was removed by performing two-10-minute washes of 25% glycerol/0.1% KOH.

## Results

### Inhibition of cholesterols synthesis reduces *axin2* expression

We have previously demonstrated that inhibition of the CSP causes craniofacial abnormalities and decreased expression of *sox10*, a marker of CNCCs (Kelsh, 2006, p. 10). However, the mechanisms by which cholesterol regulates facial development are unknown. We hypothesized that defects in cholesterol synthesis disrupt WNT signaling, consequently resulting in facial dysmorphia. To test this hypothesis, we used qPCR to monitor the expression of *axin2*, a known target gene of the WNT signaling cascade (Jho et al., 2002, p. 2) in animals carrying the Vu57 allele. As shown in Figure 1A, homozygous carriers of the Vu57 allele had a statistically significant decrease in the expression of *axin2*. We did not observe decreased expression of *axin2* in heterozygous animals. Next, we treated wildtype larvae with 1, 2, or 3uM Ro 48-8071 (4’-[6-(allylmethylamino)hexyloxy]-4-bromo-2’-fluorobenzophenone fumarate) and measured the expression of *axin2. axin2* expression was decreased at all concentrations, consistent with the prevalence of facial dysmorphia as described previously (Quintana et al., 2017) (Signore et al., 2016). These deficiencies in *axin2* expression were consistent with decreased *sox10* expression (Figure 1C).

**Figure 1:**
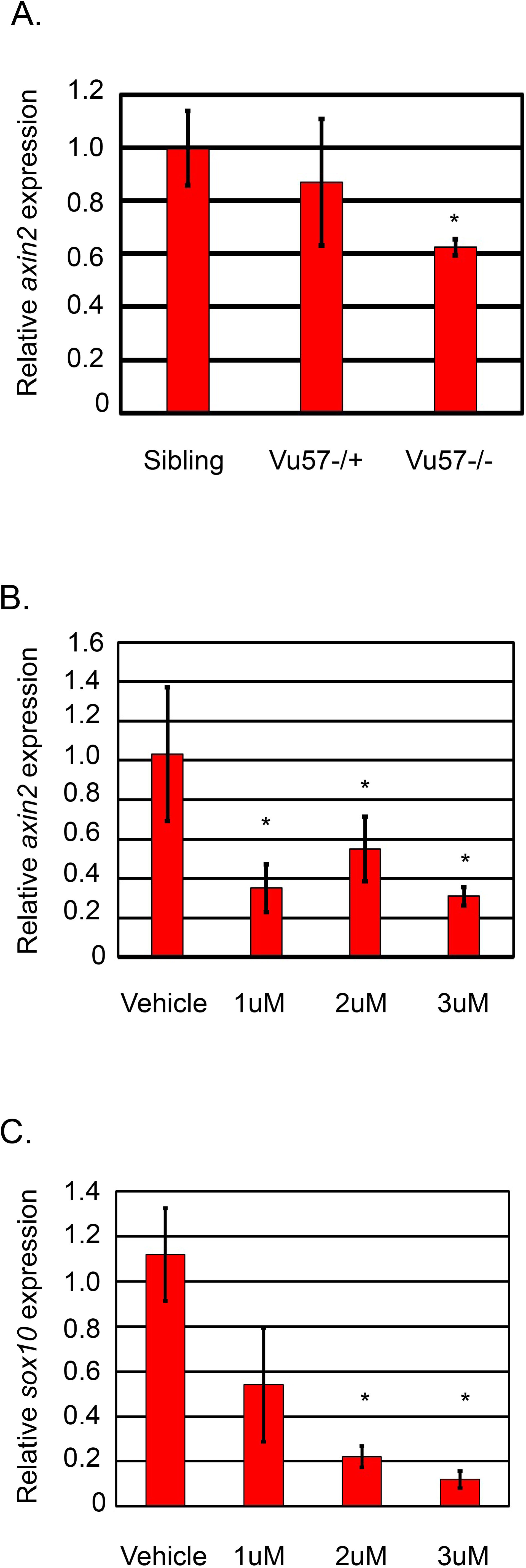
Inhibition of cholesterol synthesis is associated with decreased *axin2* expression. A. RNA was isolated from homozygous carriers of the Vu57 allele (Vu57-/-), heterozygous carriers of the Vu57 allele (Vu57+/-), or wildtype siblings (Sibling) at 24 hours post fertilization (hpf) and the relative expression of *axin2* was measured by quantitative real time PCR (qPCR). N=3/group. *P<0.05 B-C. Embryos were treated with vehicle control (DMSO), 1uM, 2uM, or 3uM Ro 48-8071 at the shield stage and total RNA was isolated at 30 hpf from a pool of embryos (N=30) and the relative expression of *axin2* (B) or *sox10* (C) was measured by qPCR. *P<0.05. Error bars represent standard deviation.

### Activation of WNT signaling restores the facial deficits in the Vu57 allele

Inhibition of the CSP is known to disrupt late stage CNCC differentiation (Quintana et al., 2017; Signore et al., 2016) and therefore, we hypothesized that activation of WNT signaling prior to the onset of these defects might restore the facial phenotypes present in the Vu57 allele. To test this, we treated offspring of the Vu57 allele with Wnt-Agonist 1. We initiated treatment at 30 hpf, removed the treatment at 54 hpf, and then assayed for craniofacial defects at 4 dpf using alcian blue. As previously described, vehicle control treated homozygous carriers of the Vu57 allele had significant craniofacial defects including the complete loss of the Meckel’s cartilage, ceratohyal, and ceratobranchial cartilages (Figure 2B). In contrast, treatment with 0.1uM Wnt-agonist 1 restored the facial defects to wildtype levels (Figure 2D; p<0.0001). Treatment with higher levels of the Wnt-agonist 1 affected survival (Table 1).

**Table 1.** Presence of facial abnormalities after treatment with WNT-Agonist I in the Vu57 allele. Craniofacial phenotypes in siblings of the hmgcs1^vu57^ allele (wildtype), heterozygous carriers, and homozygous carriers treated with vehicle control (DMSO) or different concentrations of WNT agonist-1(WNT). Phenotype indicates presence of a truncated Meckel’s cartilage, absence of an inverted ceratohyal and/or loss of the posterior ceratobranchial cartilages.

| Treatment | Wildtype (WT) | # WT Affected | Heterozygous (HET) | # HET Affected | Homozygous Mutant (MT) | # MT Affected | Total Embryos | Percent Survival (%) |
| --- | --- | --- | --- | --- | --- | --- | --- | --- |
| None | 14 | 0/14 | 25 | 6/25 | 17 | 17/17 | 56 | 100% |
| DMSO | 25 | 0/25 | 35 | 5/35 | 25 | 25/25 | 85 | 100% |
| WNT 0.1 uM | 15 | 0/15 | 32 | 7/32 | 30 | 16/30 | 77 | 100% |
| WNT 0.75 uM | 0 | 0 | 0 | 0 | 0 | 0 | 31 | 0 |
| WNT 1.0 uM | 0 | 0 | 0 | 0 | 0 | 0 | 11 | 0 |

**Figure 2:**
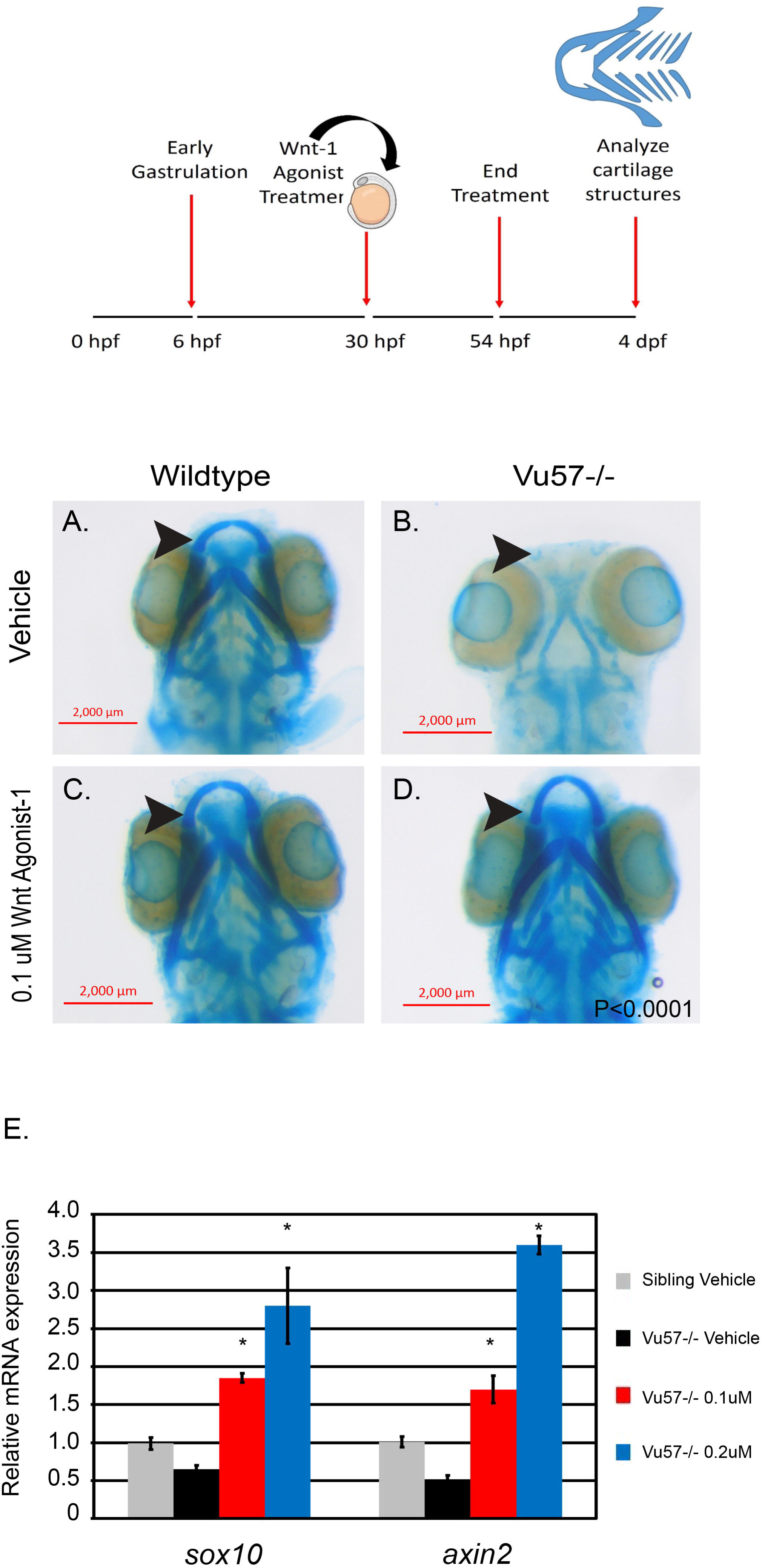
Activation of WNT signaling restores the facial defects present in the Vu57 allele. Top panel: Experimental design schematic with onset of treatment with WNT-Agonist I at 30 hours post fertilization (hpf) and removal of treatment at 54 hpf. A-D. Alcian blue staining was performed at 4 days post fertilization (dpf) in homozygous carriers of the Vu57 allele (Vu57-/-) and wildtype siblings (Sibling). Total numbers of animals reflected in Table 1. *P=0.0001. G. Homozygous carriers of the Vu57 allele (Vu57-/-), or wildtype siblings (Sibling) were treated with vehicle control (DMSO) or WNT-agonist I at either 0.1 or 0.2uM concentration. Total RNA was isolated from a pool of embryos (N=3/genotype) and the expression of *sox10* or *axin2* was measured by quantitative PCR (qPCR). Error bars represent standard deviation. *P<0.05.

We next measured the expression of *sox10* and *axin2* in Vu57 larvae treated with 0.1 or 0.2uM Wnt-agonist 1. Homozygous carriers of the Vu57 allele had a statistically significant decrease in *sox10* and *axin2* expression, consistent with craniofacial defects and decreased WNT signaling (Figure 2E). In contrast, homozygous Vu57 larvae treated with 0.1uM WNT agonist-1 had normal expression of both *sox10* and *axin2*, indicative of a restoration of the craniofacial and WNT deficits present in mutant larvae. Treatment with 0.2uM WNT agonist-1 increased the expression of *sox10* and *axin2* well above wildtype levels (Figure 2E).

### Inhibition of cholesterol reduces *axin2* expression, but inhibition of isoprenoids does not

The CSP produces cholesterol and isoprenoids and we have previously shown that inhibition of either lipid causes facial defects (Quintana et al., 2017). Both lipid products participate in cell signaling and therefore, we asked whether defects in *axin2* expression were cholesterol dependent. To test this we treated wildtype embryos with 1) Ro 48-8071, to inhibit cholesterol, 2) lonafarnib, to inhibit isoprenoids, or 3) atorvastatin, which inhibits the synthesis of both cholesterol and isoprenoids and measured the expression of *axin2* by qPCR. Treatment with either atorvastatin or Ro 48-8071 resulted in a statistically significant decrease in *axin2* expression (Figure 3A). In addition, whole mount *in situ* hybridization to detect *axin2* expression demonstrated decreased expression in the anterior region of embryos treated with Ro 48-8071 (Figure 3B&C).

**Figure 3:**
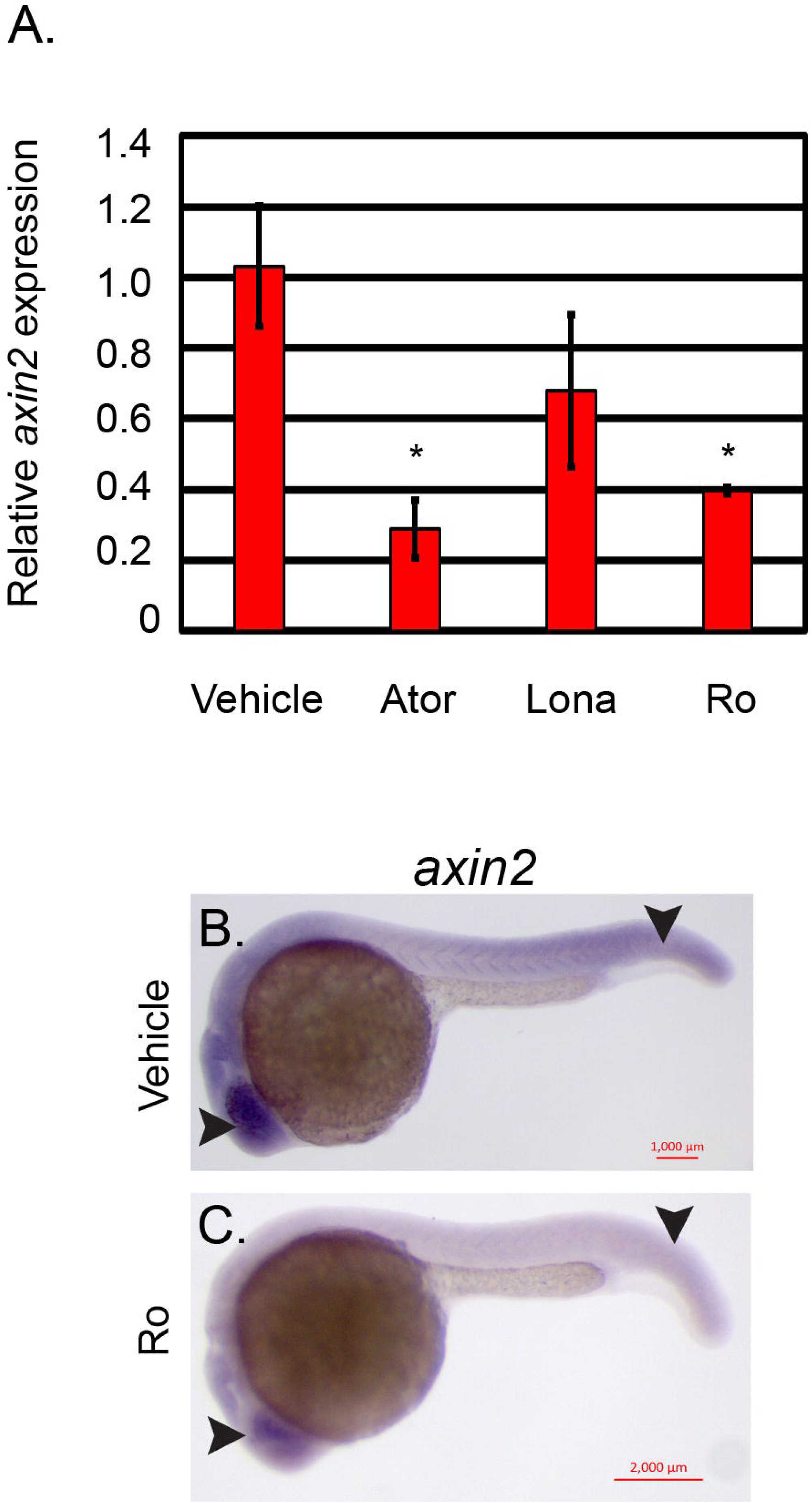
Inhibition of cholesterol, but not isoprenoids is associated with decreased *axin2*. A. Wildtype larvae were treated with vehicle control (DMSO), 2uM atorvastatin (ATOR), 8uM lonafarnib (lona), or 3uM Ro-48-8071 at shield stage. Total RNA was isolated from a pool of embryos (N=14) and the relative expression of *axin2* was measured by quantitative real time PCR (qPCR). B&C. Whole mount *in situ* hybridization was performed at 26 hours post fertilization (hpf) after treatment with 3uM R0-48-8071 or vehicle control (DMSO). Arrows indicate decreased expression in head and trunk.

## Discussion

We have previously demonstrated that mutation of the zebrafish *hmgcs1* gene (the Vu57 allele) causes craniofacial defects that are caused by defects in late stage NCC differentiation (Quintana et al., 2017). These data are supported by other zebrafish studies analyzing the craniofacial defects present upon mutation of *hmgcr* (Signore et al., 2016). Despite these advances, a molecular mechanism by which facial abnormalities develop in these animals has not been characterized. Multiple studies have suggested that defects in cholesterol synthesis are related to defects in SHH signaling (Shin et al., 2019), however homozygous mutation of the Vu57 allele does not affect SHH signaling (Quintana et al., 2017).

Cholesterol is required for the activation of various cell signaling cascades (Shimomura et al., 2019), including SHH and WNT, both of which are required for craniofacial development (Shimomura et al., 2019; Shin et al., 2019). For example, WNT activation requires the activity of disheveled, a scaffolding protein (Bernatik et al., 2011). The Disheveled protein requires cholesterol modification of its PDZ domain for proper activation (Sheng et al., 2014). Based upon these data, we hypothesized that defects in cholesterol synthesis would decrease WNT activation. Consistent with this hypothesis, we observed decreased activation of *axin2*, a WNT downstream target gene (Jho et al., 2002). Moreover, activation with a WNT agonist restored the facial phenotypes present in the Vu57 allele suggesting that the CSP regulates facial development in a WNT dependent manner.

We have previously demonstrated that isoprenoids regulate facial development in a cholesterol independent manner (Quintana et al., 2017). Isoprenoids are necessary for protein prenylation and cell signaling (Lange et al., 2000) and are known to affect several signaling pathways including WNT (Robin et al., 2014). Based upon these data, we tested whether inhibition of cholesterol or isoprenoids affected WNT signaling. We found that inhibition of cholesterol was associated with decreased *axin2*, however, inhibition of isoprenoid synthesis did not statistically reduce the expression of *axin2*. Treatment with atorvastatin caused a significant reduction in *axin2*, whereas treatment with Ro 48-8071 was more moderate. Therefore, we cannot complete rule out the effects of lonafarnib treatment on WNT signaling. However, our results suggest there are cholesterol dependent mechanisms regulating WNT activity.

Here we demonstrate that inhibition of the CSP is associated with decreased *axin2*, a downstream target gene of the WNT signaling cascade. Moreover, our work demonstrates that activation of WNT in larvae harboring the Vu57 allele completely restores the craniofacial defects present in homozygous mutants and the level of *axin2* expression in mutant larvae. These data implicate cholesterol as a key component of WNT signaling and craniofacial development.

## Acknowledgements

These studies were supported by grants from the National Institutes of Health (National Institute on Minority Health and Health Disparities-2G12MD007592 to The University of Texas El Paso, National Institutes of General Medical Sciences RL5GM118969, TL4GM118971, R25GM069621-11, UL1GM118970 to The University of Texas El Paso, and National Institute of Neurological Disorders and Stroke 1K01NS099153-01A1 to Anita M. Quintana. This study was designed, performed, and analyzed by the authors. The funding sources provided financial support for the experiments described.

## Data Sharing Statement

All data and/or model organisms associated with this manuscript are freely available upon request from the corresponding author.

